# A suppressor screen *in C. elegans* identifies a multi-protein interaction interface that stabilizes the synaptonemal complex

**DOI:** 10.1101/2023.08.21.554166

**Authors:** Lisa E. Kursel, Jesus E. Aguayo Martinez, Ofer Rog

## Abstract

Successful chromosome segregation into gametes depends on tightly-regulated interactions between the parental chromosomes. During meiosis, chromosomes are aligned end-to-end by an interface called the synaptonemal complex, which also regulates exchanges between them. However, despite the functional and ultrastructural conservation of this essential interface, how protein-protein interactions within the synaptonemal complex regulate chromosomal interactions remains poorly understood. Here we describe a novel interaction interface in the *C. elegans* synaptonemal complex, comprised of short segments of three proteins, SYP-1, SYP-3 and SYP-4. We identified the interface through a saturated suppressor screen of a mutant that destabilizes the synaptonemal complex. The specificity and tight distribution of suppressors point to a charge-based interface that promotes interactions between synaptonemal complex subunits and, in turn, allows intimate interactions between chromosomes. Our work highlights the power of genetic studies to illuminate the mechanisms that underly meiotic chromosome interactions.

**Significance Statement:** Gamete production requires tightly regulated interactions between the parental chromosomes, which co-align and exchange information. These events are mediated by the synaptonemal complex – a thread-like structure that assembles between the parental chromosomes. The molecular interactions that underly synaptonemal complex assembly remain poorly understood due to rapid sequence divergence and challenges in biochemical reconstitution. Here we identify a novel three-component interface in the nematode synaptonemal complex. Destabilization and subsequent restoration of this interface link the integrity of the synaptonemal complex with chromosome alignment and regulation of exchanges. Beyond mechanistic understanding of chromosomal interactions, our work provides a blueprint for genetic probing of large cellular assemblies that are refractory to structural analysis and sheds light on the forces that shape their evolution.

## Introduction

Sexual reproduction requires tightly regulated chromosomal interactions. During meiosis, parental chromosomes (homologs) pair and align along their length. Chromosome alignment sets the stage for reciprocal exchanges called crossovers that physically link the homologs and allow them to correctly segregate during the first meiotic division. Failure to form crossovers, or formation of too many crossovers, can lead to chromosome missegregation, aneuploid progeny and infertility (1–3).

Crossovers form within, and are regulated by, a dedicated meiotic chromosomal structure called the synaptonemal complex (SC). Upon entry into meiosis, each homolog adopts an elongated morphology organized around a backbone called the axis. The axes of the two homologs are then aligned, end-to-end, concomitant with the assembly of the SC. While the axes are classically considered part of the SC, here we use the term SC to specifically refer to the structure assembled between parallel axes, also known as the central region of the SC. In addition to bringing the homologs, and therefore the exchanged DNA sequences that form crossovers, into proximity, the SC directly participates in regulating crossover number and distribution (4, 5).

Our understanding of how the SC brings homologs together and regulates crossovers is limited. While the ultrastructure of the SC is conserved across eukaryotes, appearing as a ∼100nm-wide ladder in electron-micrographs (6–11), the protein sequences of its components diverge rapidly (12–14). The poor sequence homology has slowed the identification of SC components, and it is unclear whether the complete set of components has been defined in any organism. Poor sequence homology has also limited the translation of structural findings across model organisms. Finally, we have very limited molecular-genetic information since only a small subset of components has been subjected to systematic mutational analysis or biochemical and structural probing.

The nematode *C. elegans* has proved a useful model for studying SC structure and function. The SC in worms assembles prior to crossover formation and is essential for all crossovers (8). In addition, the role of the SC in regulating crossover distribution has been directly demonstrated (15). The worm SC includes eight proteins: SYP-1, -2, -3, -4, -5/6 and SKR-1/2 (SYP-5 is partially redundant with SYP-6 and SKR-1 is partially redundant with SKR-2; (8, 16–21)). These components exhibit stereotypical organization within the SC: SYP-1 and -5/6 are organized in a head-to-head fashion to span the distance between the axes, with their N-termini in the middle of the SC and C-termini near the axis. The other components localize to a band in the middle of the SC (Figure 1A, (16, 22, 23)).

**Figure 1:**
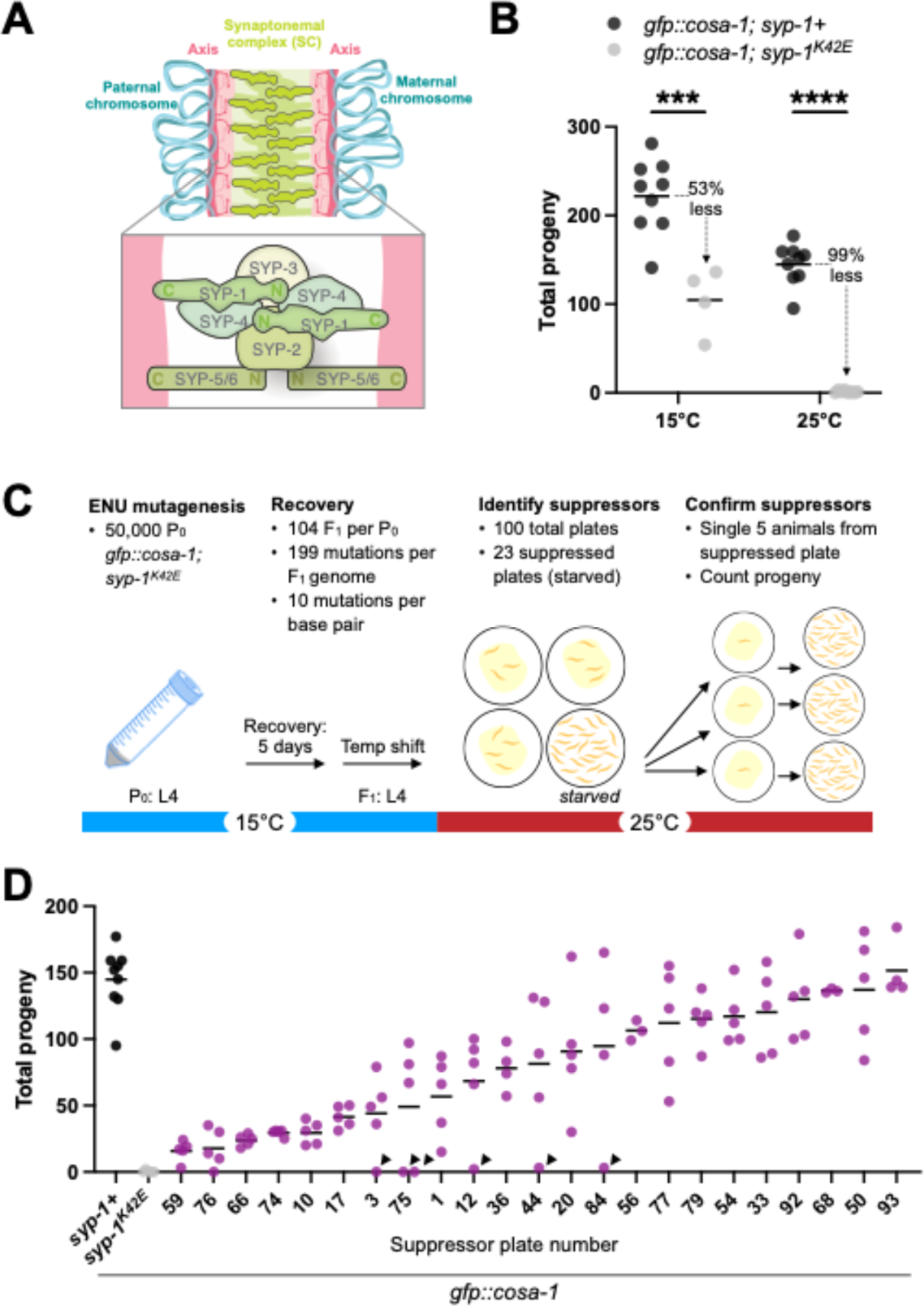
Mutagenesis screen identifies suppressors of *syp-1^K42E^* fertility defect. (A) Top, cartoon of paired homologous chromosomes (blue) with the synaptonemal complex (SC, green) assembled between them. Bottom, cartoon depicting six SC proteins from *C. elegans* with the N- and C-termini of SYP-1 and SYP-5/6 labeled. (B) Total self-progeny from *syp-1+* and *syp-1^K42E^* animals in a *gfp::cosa-1* background at 15°C and 25°C. Asterisks indicate statistical significance of the comparison of total progeny between *gfp::cosa-1; syp-1+* and *gfp::cosa-1; syp-1^K42E^* at 15°C and 25°C using an unpaired t-test. (C) Schematic of suppressor screen. We mutagenized 50,000 *gfp::cosa-1; syp-1^K42E^*animals (P^0^) using ENU and allowed them to recover at 15°C. After five days, when most F^1^s were at the L4 developmental stage, we shifted the worms to 25°C to apply selection. Suppressed plates were able to consume all of the bacterial lawn and starve the plate. We confirmed the suppression phenotype by picking five single worms from each suppressed plate and counting self-progeny at 25°C (quantified in D). (D) Total self-progeny from worms singled from 23 suppressed plates sorted by degree of suppression. Arrowheads indicate plates with zero to three total self-progeny. These plates likely contained non-suppressed animals, suggesting that when the worms were singled the selective sweep by the suppressors was still incomplete. In (A) and (C), mean values are shown by a horizontal black line and asterisks indicate p-values as follows: *** < 0.001, **** < 0.0001.

Previously, we carried out mutational analysis of the N-terminus of SYP-1 and demonstrated its role in holding the parallel axes together and in regulating crossovers (24). Here, we use a temperature-sensitive single substitution mutant in *syp-1* to carry out a suppressor screen that allowed us to define a novel interaction interface in the middle of the SC. This interface ensures efficient assembly of the SC onto chromosomes, and consequently, the formation of tightly regulated crossovers and successful chromosome segregation. Many of the suppressors we isolated did not affect meiosis by themselves despite altering highly conserved residues, illuminating the unusual selective pressures that shape SC protein evolution.

## Results

### A suppressor screen of a temperature-sensitive SC mutant

We previously identified a lysine-to-glutamic acid substitution, *syp-1^K42E^,* that weakens interactions among SC proteins, causing failures in chromosome synapsis and crossover regulation (24). In *syp-1^K42E^,* the stability of the SC progressively weakens as the temperature increases, leading to worsening meiotic phenotypes (24). At 15°C, the SC in *syp-1^K42E^* animals is mostly unaffected, and the worms indeed produce many viable self-progeny, although their number is reduced by 53% compared to control animals (Figure 1B, mean total progeny 15°C; *syp-1+* = 222, *syp-1^K42E^* = 104). At 25°C, a temperature at which wild-type *C. elegans* can reproduce without discernable meiotic defects, *syp-1^K42E^* animals are nearly sterile (Figure 1B, mean total progeny 25°C; *syp-1+* = 145, *syp-1^K42E^* = 1). The SC at this temperature assembles onto unpaired chromosomes, a morphology that is never observed in wild-type animals (24).

In order to delineate the dimerizing SYP-1 interface, we carried out a whole-genome mutagenic screen to identify suppressors of the *syp-1^K42E^*fertility defect. We used the alkylating agent *N*-ethyl-*N*-nitrosourea (ENU) to mutagenize ∼50,000 P^0^ *gfp::cosa-1; syp-1^K42E^* animals grown at 15°C and divided them among 100 plates. We allowed their F^1^ progeny (carrying heterozygous mutations) to grow at 15°C until they reached the young adult (L4) stage, at which point we shifted them to the non-permissive temperature of 25°C. Candidate suppressors of the *syp-1^K42E^* fertility defect quickly outcompeted their non-suppressed siblings and starved the plate (Figure 1C). Through this approach, we identified 23 suppressed lines that showed a wide range in their ability to suppress the infertility of *syp-1^K42E^* at 25°C (Figure 1D).

### All suppressed lines of syp-1^K42E^ contain mutations in SC proteins

To identify the causative suppressor mutations, we employed a combination of candidate Sanger sequencing and whole genome sequencing followed by variant calling (Figure 2A). We hypothesized that suppressor mutations could be within SC components, so we sequenced the N-terminus of *syp-1* and *syp-3*. (SYP-3 also localizes in the middle of the SC (22), and a small truncation in *syp-3* exhibits phenotypes similar to those of *syp-1^K42E^*(18)). This approach had the added advantage of confirming the presence of the original *syp-1^K42E^* mutation. Twenty-two out of the 23 suppressed strains had the original *syp-1^K42E^* mutation. The remaining strain had acquired a mutation that resulted in reversion to the wild-type SYP-1 sequence (Figure 2A, *syp-1^E42K^*revertant). We could distinguish the revertant from wild-type contamination because the suppressed strain used AAA to encode the lysine (K) at position 42, rather than the AAG codon in the reference genome (Figure 2B). We also identified seven additional candidate suppressor mutations in *syp-1* and three candidate suppressor mutations in *syp-3.* In *syp-1,* there were three charge-altering mutations*, syp-1^E41K^* and *syp-1^E45K^*(recovered twice independently), three polar to hydrophobic mutations, *syp-1^S24L^* (twice) and *syp-1^T35I^,* and one hydrophobic to hydrophobic mutation, *syp-1^V39I^.* The three mutations in *syp-3* were at amino acid position 62 and altered the normally negatively charged aspartic acid (D) to either asparagine (N, polar), valine (V, hydrophobic) or glycine (G, small/hydrophobic).

**Figure 2:**
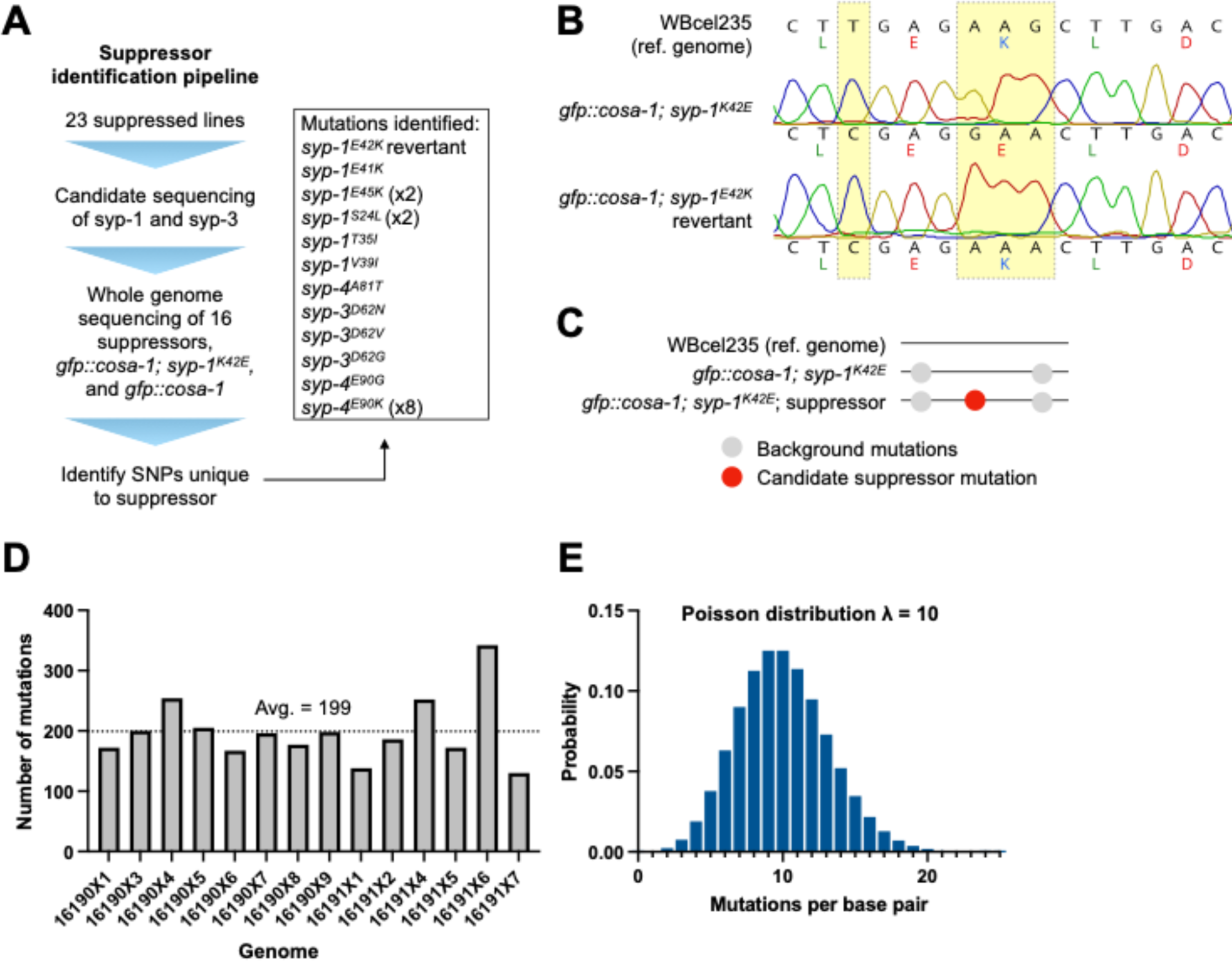
All suppressor strains contain mutations in SC proteins. (A) Pipeline used to identify suppressor mutations. We performed Sanger sequencing on portions of *syp-1* and *syp-3* in all suppressed lines to confirm the presence of the original *syp-1^K42E^*mutation and to identify possible suppressor mutations. We performed whole genome sequencing on 16 suppressor strains and controls. We used GATK to call variants unique to each suppressor strain and found an additional 14 unique mutations in in SC proteins, some of which occurred multiple times independently. (B) Chromatograph traces from Sanger sequencing of *syp-1* in *gfp::cosa-1; syp-1^K42E^* and the revertant strain identified in the screen (*gfp::cosa-1; syp-1^E42K^*) aligned to the reference genome. The reference genome and revertant strain each encode the lysine (K) at position 42 (yellow box) with a different codon (AAG and AAA, respectively). In addition, the silent T->C mutation in the codon encoding a lysine at position 40, which was introduced while creating the original *syp-1^K42E^* mutation, is present in the revertant. (C) Schematic depicting the strategy used for variant calling. We identified SNPs and indels in *gfp::cosa-1; syp-1^K42E^* and in the suppressed lines. Mutations present in *gfp::cosa-1; syp-1^K42E^* but not in the reference genome were considered background mutations. We subtracted these background mutations from the mutations in each suppressed strain to generate a list of putative suppressor mutations. (D) Bar graph showing the number of unique, non-background mutations in each sequenced suppressed line. Average number of mutations per genome = 199. (E) Histogram of Poisson distribution for λ = 10, the average number of ENU-generated mutations per base pair in the screen. The likelihood that substitution in a base pair was not screened for suppression is < 5x10^-5^.

We identified the remaining suppressor mutations through whole genome sequencing. We identified all homozygous SNPs and indels in the suppressed strains relative to the parental strain and the reference genome, WBcel235 (Figure 2C). In addition to the 2,036 SNPs present in *gfp::cosa-1* relative to the reference genome, each suppressed genome had 199 homozygous mutations on average (Figure 2D). Using this number, plus the number of mutagenized P^0^ animals (50,000), the number of F^1^s per P^0^ at 15°C (Figure 1A, *gfp::cosa-1; syp-1^K42E^* average total progeny = 104), and the haploid genome size of *C. elegans* (1 x 10^8^ base pairs), we estimate that each base pair in the genome was mutagenized 10 times in our screen. Using a Poisson distribution, the probability that a base pair was not screened is 4.5x10^-5^ (Figure 2E, λ=10).

Out of the 2,789 homozygous mutations in the 14 independent suppressed lines we sequenced, 1,968 were in intergenic regions and UTRs, 159 were synonymous site mutations and 96 mutations were in gene introns (Figure S1A, Table S1), in line with the genomic distribution of these elements. We found 537 missense mutations and 29 mutations that were likely to abolish gene function through the loss of a start or a stop codon, a nonsense or frameshift mutation or a mutation of a splice acceptor/donor site (Figure S1A). The distribution of SNPs matched the expected mutagenic profile of ENU with an overall bias towards GC > TA transitions, although all transitions and transversions were represented (Figure S1B, (25)).

To prioritize suppressor mutations, we sorted for genes whose RNAi phenotype, allele phenotype or gene ontology included the words “meiosis” or “meiotic” (Table S2). Each genome had between one and four missense mutations in meiotic genes. Strikingly, every genome had a missense mutation in an SC subunit. These included three mutations in *syp-1* (*syp-1^S24L^*[identified twice], *syp-1^T35I^* and *syp-1^V39I^*) and three mutations in *syp-4* (*syp-4^A81T^, syp-4^E90G^*and *syp-4^E90K^* [identified eight times]).

The location of the suppressors in a narrow region of the same SC proteins and the independent isolation of multiple instances of the same mutation, like *syp-4^E90K^,* suggests that the mutations in SYP-1, -3 and -4 are the causative suppressor mutations.

### Suppression strength correlates with amino acid charge

We noticed that many of the suppressors change a negatively charged residue to a positively charged residue (*e.g., syp-1^E41K^, syp-1^E45K^, syp-4^E90K^*) or neutralize a negatively charged residue (*e.g., syp-3^D62N^, syp-3^D62V^, syp-3^D62G^, syp-4^E90G^*). Accordingly, we divided the suppressors into three categories and carefully analyzed their effects; one, suppressor mutations that neutralized a negative charge (*syp-3^D62N^, syp-3^D62V^*and *syp-4^E90K^),* two, suppressor mutations that altered a polar residue (*syp-1^T35I^* and *syp-1^S24L^*), and three, suppressor mutations that altered a hydrophobic residue (*syp-4^A81T^*and *syp-1^V39I^*). Rather than solely rely on homozygous suppressed animals recovered from the screen, we engineered three of the strongest suppressor mutations (*syp-3^D62N^, syp-3^D62V^* and *syp-4^E90K^*) using CRISPR/Cas9 in the *gfp::cosa-1; syp-1^K42E^* background. The suppression we observed in these engineered strains (see below) rules out a significant causative role for non-SC mutations and further confirms the identity of the suppressors in SYP-3 and -4.

The original mutation, *syp-1^K42E^,* has very low total progeny and a high incidence of male self-progeny at 25°C (Figure 1B and (24)), caused by meiotic nondisjunction of the X chromosomes. Animals suppressed with *syp-3^D62N^, syp-3^D62V^*or *syp-4^E90K^* showed complete suppression of defects in total progeny and percent males at 25°C (Figure 3A, 3B). Like wild-type animals, they had large brood sizes (Figure 3A, blue-filled circles, *gfp::cosa-1; syp-1^K42E^; syp-3^D62N^* = 223, *gfp::cosa-1; syp-1^K42E^; syp-3^D62V^* = 210, *gfp::cosa-1; syp-1^K42E^; syp-4^E90K^* = 189) and rare male self-progeny (Figure 3B, blue-filled circles, all strains <1% male progeny). Animals suppressed with *syp-1^T35I^* and *syp-1^S24L^* showed an intermediate phenotype with average total progeny of 178 and 99, respectively, and a slightly elevated percentage of male self-progeny (Figure 3A, 3B, teal-filled circles, *gfp::cosa-1; syp-1^K42E+T35I^* = 1.4% and *gfp::cosa-1; syp-1^K42E+S24L^* = 4% male progeny). Finally, animals suppressed with *syp-4^A81T^* and *syp-1^V39I^* provided the weakest suppression and had relatively low total progeny and a high percentage of male offspring, albeit significantly suppressed compared to *gfp::cosa-1; syp-1^K42E^* (Figure 3A, 3B, yellow filled circles, *gfp::cosa-1; syp-1^K42E^; syp-4^A81T^*average total progeny = 25, average percent males = 9.5%, *gfp::cosa-1; syp-1^K42E+V39I^* average total progeny = 31, average percent males = 14.8%).

**Figure 3:**
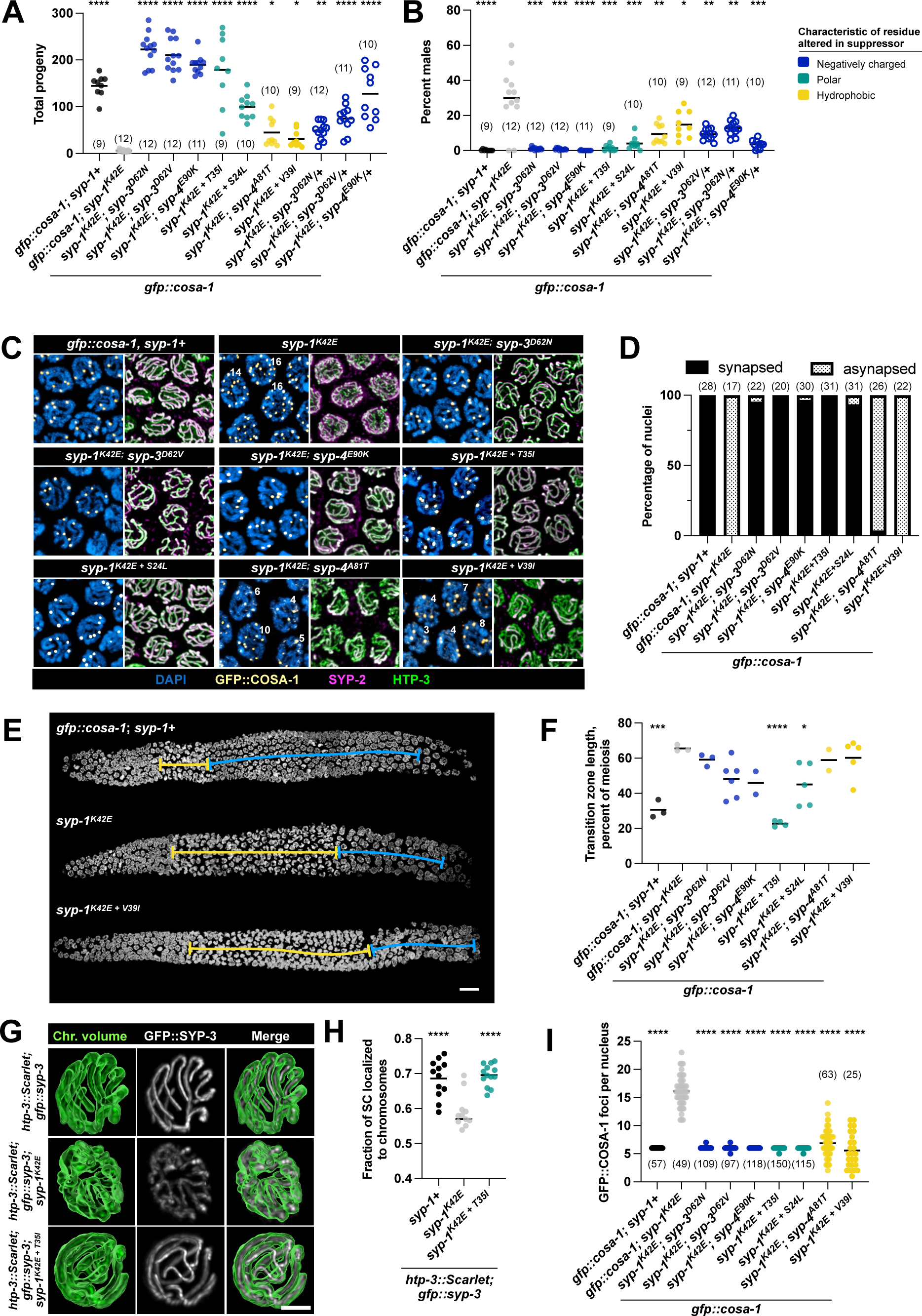
Strength of suppression correlates with amino acid charge. Total progeny (A), and percent male self-progeny (B), for seven representative suppressors at 25°C. Phenotype of the heterozygous state of the three strongest suppressors is also shown in (A) and (B). Number of replicates are listed in parentheses. Throughout the figure, suppressor strains are colored according to the characteristic of the residue that was mutated in the screen. (C) Immunofluorescence images of pachytene nuclei showing the extent of synapsis and GFP::COSA-1 foci for representative suppressed genotypes at 25°C. The number of foci is indicated in nuclei with more or less than the six expected GFP::COSA-1 foci. Scale bar = 5 µm. (D) Bar graphs showing percentage of early pachytene nuclei with partially or completely asynapsed chromosomes at 25°C. N values in parentheses indicate the number of nuclei examined. (E) Example images of whole gonads stained with DAPI used to measure transition zone length. Transition zones are marked with a yellow line, pachytene is marked with a blue line. Scale bar = 10 µm. (F) Quantification of transition zone length as percent of meiosis at 25°C. (G) Images of chromosome volumes generated using HTP-3::wrmScarlet fluorescence signal and live fluorescence images of GFP::SYP-3 taken at 20°C. (H) Dot plot showing percent SC signal localized to chromosomes relative to total nucleoplasmic fluorescence in a single nucleus at 20°C. (I) Number of GFP::COSA-1 foci per nucleus in suppressed strains at 25°C. Note that only the two weak suppressors, *syp-1^K42E^; syp-4^A81T^* and *syp-1^K42E^ ^+ V39I^,* have an elevated standard deviation (stdev = 2.38 and 3.11, respectively) compared with *syp-1+* (stdev = 0). For panels (A – C), (H) and (I), asterisks (black) indicate statistical significance compared to *gfp::cosa-1; syp-1^K42E^* and horizontal black lines show mean values. Asterisks are representative of p-values as follows: * < 0.05, ** < 0.01, *** < 0.001, **** < 0.0001. See methods for detailed description of statistical analyses.

We hypothesized that if the suppressor sites in SYP-3 and -4 physically interact with SYP-1^K42E^, then even one copy of a suppressor mutation may allow the suppressor proteins to assemble into the SC and restore interactions among some of the SC proteins. We therefore also tested three of the strongest suppressors and found that heterozygous *syp-3^D62N^, syp-3^D62V^*and *syp-4^E90K^* provided an intermediate suppression phenotype (Figure 3A, 3B, blue open circles). This also implies that the selection in our screen began in the F^1^ rather than the F^2^ generation since the F^1^s could have benefited from a heterozygous suppressor mutation when they were shifted to the nonpermissive temperature (Figure 1C).

### Suppressor mutations restore SC assembly, SC stability and crossover regulation

We next verified that the suppressors restored the SC defects in *syp-1^K42E^*animals. At 20°C, *syp-1^K42E^* nuclei in mid-pachytene fail to synapse homologs end-to-end and typically have at least one fully asynapsed and one partially synapsed (forked) chromosome pair; at 25°C, the SC assembles onto unpaired chromosomes (24). We found that strains suppressed by *syp-3^D62N^, syp-3^D62V^, syp-4^E90K^, syp-1^T35I^*and *syp-1^S24L^* achieved complete synapsis by mid-pachytene at 25°C (Figure 3C, 3D). We saw almost no evidence of forked chromosomes or unpaired chromosomes lacking SC (Figure 3C, 3D, Figure S2). In contrast, strains suppressed by *syp-4^A81T^* and *syp-1^V39I^* failed to achieve full synapsis and only had a few synapsed chromosomes per nucleus (Figure 3C, 3D, Figure S2), likely accounting for their failure to form crossovers between all homolog pairs, and, consequently, their smaller brood sizes and high percentage of male self-progeny.

Previous work has shown that meiotic nuclei sense unpaired chromosomes, and, in response, spend a longer time in the so-called transition zone, where homology search and SC assembly occur (corresponding to the classically defined leptotene and zygotene stages of meiosis; (26)). The extended transition zone could allow for more time to complete SC assembly. We found that most suppressed strains failed to suppress the extended transition zone of *syp-1^K42E^* animals (Figure 3E-F, Figure S3). In the case of the weak suppressors, that is likely due to partial synapsis (Figure 3C-D). Interestingly, the strong and intermediate suppressors exhibited different transition zone lengths – extended in the strong suppressors and shorter in the intermediate suppressors – even though almost all nuclei in these strains achieved complete synapsis by mid-pachytene (Figure 3C-D). The mechanism that senses and responds to asynapsed chromosomes depends on axis proteins, likely involving exposed axes not associated with the SC (27, 28). A possible reason for the difference in transition zone length between the strong and intermediate suppressors might be their different propensities to associate with the axes, either in the context of assembled SC or with aberrant conformations, such as forked chromosomes. Alternatively, the difference between the strong and intermediate suppressors may reflect slower kinetics of synapsis or defects that cannot be discerned at the resolution of confocal microscopy.

We also tested whether the suppressor mutations restore the unstable SC in *syp-1^K42E^* animals (24). We measured the proportion of GFP::SYP-3 on chromosomes relative to the total nucleoplasmic amount in *syp-1^K42E^* and suppressed (*syp-1^T35I^ ^+ K42E^*) animals. We identified inter-chromosomal regions in live animals using HTP-3::wrmScarlet, a component of the axis that assembles independently of the SC. Consistent with our published findings, we found that less GFP::SYP-3 is recruited to chromosomes in *syp-1^K42E^* compared to a wild-type control (Figure 3G-H and (24)). The suppressor mutation (*syp-1^T35I^*) restored the amount of SC recruited to chromosomes to wild-type levels (Figure 3G-H). Notably, since we analyzed animals at the semi-permissive temperature of 20°C, where mostly normal synapsis occurs in all tested strains, we likely underestimated the full destabilizing effect of *syp-1^K42E^*.

Finally, we assessed the ability of the suppressed strains to carry out accurate crossover regulation. *syp-1^K42E^*exhibits weaker crossover interference, with more than one crossover observed on the same chromosome and the same stretch of SC (24). We leveraged a cytological marker of crossovers, GFP::COSA-1, which localizes to each of the designated crossovers, numbering six in wildtype *C. elegans* (one per chromosome) (29). Mutations that impact the integrity of the SC, including *syp-1^K42E^* (24), *syp-1* RNAi (15), and *syp-4^CmutFlag^(ie25)* (22), reduce crossover interference so that there are more than six crossovers per nucleus. We found that strains suppressed by *syp-3^D62N^, syp-3^D62V^, syp-4^E90K^, syp-1^T35I^* and *syp-1^S24L^*all had six crossovers per nucleus (Figure 3C, 3I), indicating restoration of crossover interference. In contrast, the weak suppressors *syp-4^A81T^* and *syp-1^V39I^* showed a wide distribution of crossover number, with some nuclei having only one or two crossovers and some having more than 10 (Figure 3C, 3I). This is likely not just a reflection of the incomplete synapsis in these mutants, since both harbored instances of chromosomes (*i.e.*, a stretch of SC) with more than one crossover (Figure S4).

In summary, the suppressors of the fertility defect of *syp-1^K42E^*also suppressed its other meiotic phenotypes: incomplete synapsis, unstable SC and weakened crossover interference. Furthermore, weaker suppressors consistently exhibited incomplete suppression. Our data thus suggests that SYP-1 physically interacts with SYP-3 and SYP-4 via charge-charge interactions between the N-terminus of SYP-1, position 62 in SYP-3 and position 90 in SYP-4. SYP-1^K42E^ weakens this interaction and, as a result, the SC is less stable and fails to fully assemble onto chromosomes and to confer robust crossover interference (24). The suppressors restore this charge interface, and consequently the chromosomal and meiotic defects.

### Suppressors alter conserved residues in SC proteins

We next wished to address the molecular mechanism of suppression. Unfortunately, atomic structures for *C. elegans* SC proteins have not been solved, in a complex or in isolation, preventing us from localizing the mutated sites relative to one another (beyond their co-localization in the middle of the SC (22, 23)). It has also been challenging to derive structural insight from other model organisms through modeling, due to limited sequence homology between clades (12–14). Our attempts to model a docking interface between the regions surrounding the mutations using AlphaFold did not yield a high-confidence interface.

Lacking tertiary structural information, we examined the potential impact of the suppressor mutations on coiled-coil domains – a conserved feature of SC proteins throughout eukaryotes (12). The suppressor mutations in SYP-1 occur before the start of the coiled-coil domain (Figure 4A-B). In SYP-3 and SYP-4, the suppressor mutations lie within extended coiled-coil domains but do not significantly alter their predicted structure (Figure 4C-F, Figure S5A-B). In SYP-3, the site of the suppressor mutation is not contained within a heptad repeat – the distinguishing feature of coiled-coils. So, even though one of the mutations introduces a glycine, which can disrupt helices (30), the suppressors are not likely to impact the overall coiled-coil structure of the protein (Figure 4C-D, Figure S5A). In SYP-4, the suppressor mutations are found within perfect heptad repeats, where the ‘a’ and ‘d’ (first and fourth) positions are occupied by hydrophobic residues, and the ‘e’ and ‘g’ (fifth and seventh) positions are polar. The suppressor mutations do not change the underlying heptad structure: they either alter a residue in the non-significant ‘c’ position (A->T) or swap one polar residue for another in the ‘e’ position (E->K; Figure 4E-F, Figure S5B).

**Figure 4:**
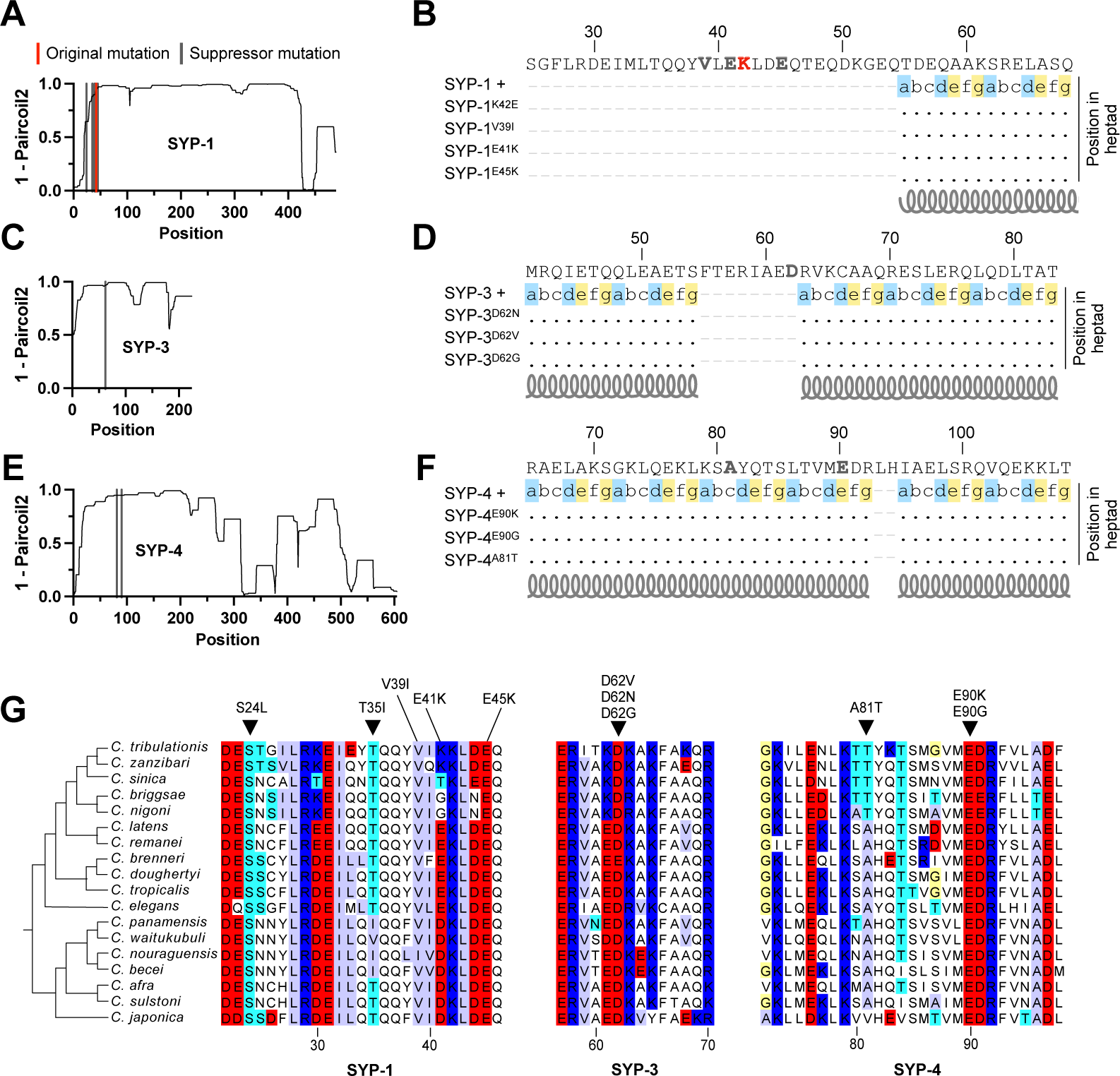
Suppressors alter conserved, charged residues in SYP-1, -3 and -4, but do not disrupt coiled-coils. (A) Coiled-coil scores per position for SYP-1. The location of original and suppressor mutation is indicated by vertical red and gray lines, respectively. A higher score implies that the region is more likely to form a coiled-coil. (B) Impact of suppressor mutations on the heptad repeats that underly coiled-coils in SYP-1. Continuous heptad repeats are indicated by their positional nomenclature, a – g, and by a grey coil. Important positions in the heptad repeats are highlighted. Position a and d (blue) are typically hydrophobic, position e and g (yellow) are typically charged or polar. Note: in SYP-1, the original and suppressor mutations occur outside the coiled-coil domain. The heptad structure is shown for SYP-1+ and is aligned to SYP-1 mutants. Dots indicate the position has the same heptad structure in the mutants. (C – F), same as (A) and (B) except for SYP-3 and SYP-4. In (B), (D), and (F), the sequence shown is from the wild-type protein and suppressor and original mutations are colored in bold gray or red letters, respectively. (G) Partial alignment of SYP-1, -3 and -4 from 18 *Caenorhabditis* species. Suppressor mutations are indicated above the alignment. Amino acids are highlighted accordingly: negatively charged = red, positively charged = blue, serine or threonine (polar) = cyan, valine, isoleucine or alanine (hydrophobic) = lilac, glycine = yellow. Note that suppressor residues are either completely conserved (*e.g.,* SYP-1^S24^ SYP-3^D62^ and SYP-4^E90^) or have been sampled by evolution (*e.g.,* SYP-1^T35^ – *C. panamensis, C. nouraguensis* and *C. becei* all encode isoleucine (I) at position 35).

The assembly of the SC through condensation (31) suggests a potential function for intrinsically disordered regions that can drive phase-separation (32). However, the mutations do not lie within conserved intrinsically disordered regions (12), arguing that the suppressors do not act by altering these domains.

To obtain hints as to the functional importance of the residues mutated in the suppressors, we analyzed their conservation in an alignment from 18 *Caenorhabditis* species, estimated to share a common ancestor ∼20 million years ago (Figure 4G, (33)). The suppressors are significantly enriched for positions that are completely conserved in *Caenorhabditis*, like the original *syp-1^K42E^* mutation (Figure 4G, *e.g., syp-1^S24L^, syp-1^E45K^, syp-4^E90K^,* chi-square test, *X^2^* (1, *N* = 22) = 15.65, p < .0001)). Whereas 16/22 suppressor mutations altered completely conserved residues, only 5% of the amino acids in SYP-1, 19% of the amino acids in SYP-3 and 16% of amino acids in SYP-4 are completely conserved. Interestingly, in many cases where the residue was not completely conserved, evolution has previously sampled the suppressor mutation: e.g., the glutamic acid to lysine substitution in position 41 in SYP-1 is also present in *C. tribulationis* and *C. zanzibari* (Figure 4G; see also *syp-1^T35I^, syp-1^V39I^ and syp-4^A81T^*). Many of the positively charged and polar residues mutated in the suppressors were conserved throughout *Caenorhabditis*, although two were not – threonine 35 and glutamic acid 41 in SYP-1. Notably, the hydrophobic residues mutated in the suppressors were both not conserved.

Consistent with the conservation of the mutated sites, we found almost no variation at suppressor sites among 550 wild isolates of *C. elegans* (34) and no variation at homologous sites in 14 wild isolates of *C. remanei* (35) (Table S3, Data S1 – S3). In *C. elegans* wild isolates, only two substitutions overlapped with suppressed sites, SYP-1^V39A^ and SYP-1^E45D^, both of which are conservative replacements that do not alter charge. Together, these findings suggest that the SYP-1/-3/-4 interface has co-evolved to maintain interactions among SC proteins.

### Suppressor mutations alone exhibit no meiotic phenotype

We next investigated the effects of the suppressor mutations alone. Since the change from a positive to a negative charge in *syp-1^K42E^*causes severe synapsis and crossover regulation defects, we hypothesized that the charge-reversing suppressors would exhibit similar phenotypes by themselves. We analyzed the three strongest suppressors, *syp-3^D62N^*, *syp-3^D62V^* and *syp-4^E90K^*as well as a strain that combined two suppressor mutations, *syp-3^D62V^; syp-4^E90K^*. In most conditions, there was not a significant decrease in the number of progeny or increase in the percentage of male self-progeny in the suppressors when compared to the *gfp::cosa-1* control at 15°C, 20°C and 25°C (Figure 5A-C). The only exception was the double suppressor, *syp-3^D62V^; syp-4^E90K^,* which demonstrated a mild (31%) but statistically significant reduction in total progeny at 25°C (Figure 5C). We also quantified crossover events and asynapsis and found no evidence of defects in either *syp-3^D62N^, syp-3^D62V^, syp-4^E90K^* or *syp-3^D62V^; syp-4^E90K^* (Figure 5D-F).

**Figure 5.**
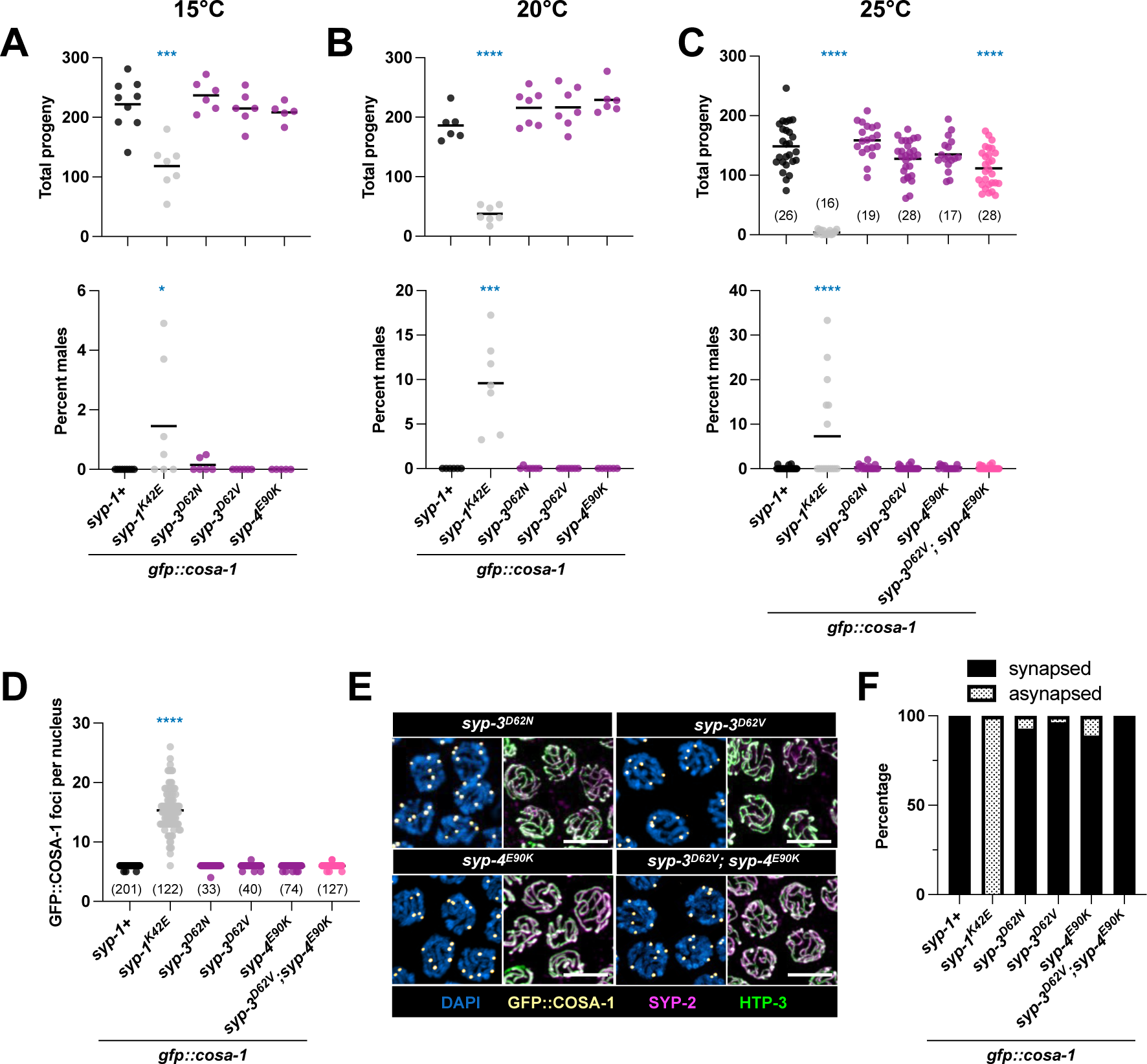
Suppressors alone do not affect the SC. Total progeny and percent male progeny for suppressors alone at 15°C (A), 20°C (B) and 25°C (C). Note that (C) also contains a strain with two suppressor mutations combined (*gfp::cosa-1; syp-3^D62V^, syp-4^E90K^*). We used an ordinary one-way ANOVA with Dunnett’s test for multiple comparison to test for a difference of means for total progeny (A – C, top), and we use a Kruskal-Wallis test with Dunn’s test for multiple comparison to test for differences in percent males among genotypes (A – C, bottom). (D) Number of GFP::COSA-1 foci per nucleus. (E) Immunofluorescence images of mid-pachytene nuclei showing GFP::COSA-1 foci and extent of synapsis. (F) Bar graphs showing percentage of early pachytene nuclei with partially or completely asynapsed chromosomes. For panels (A – D), asterisks (blue) indicate statistical significance compared to *gfp::cosa-1* and horizontal black lines indicate mean values. Asterisks are representative of p-values as follows: * < 0.05, ** < 0.01, *** < 0.001, **** < 0.0001. See methods for detailed description of statistical analyses.

While the single mutants do not appreciably affect meiosis at typical growth conditions, we wondered whether they might affect the response to extreme environments. Meiosis in general, and the SC in particular, are highly temperature-sensitive, even in wild-type plants and animals (36, 37). In *C. elegans*, a shift in temperature from 25°C to 26.5°C causes SC aggregation in early pachytene, although SC that is already assembled seems protected from this fate (36). We found no difference between *syp-4^E90K^; gfp::cosa-1* and the *gfp::cosa-1* control in their response to elevated temperature (Figure S6). In both strains, there were SC aggregates in early pachytene but not late pachytene after a 16-hour exposure to 26.5°C, suggesting that the SC in *gfp::cosa-1; syp-4^E90K^* is neither sensitized nor resistant to high temperatures.

## Discussion

The molecular interfaces that allow the SC to mediate interactions between the homologs remain a major gap in our understanding of meiotic chromosome dynamics. The identity and nature of these interfaces are poorly understood, largely because the SC has proven a difficult candidate for traditional structural approaches. An exception is the mammalian SC, where multiple recombinant SC proteins have been subjected to structural analysis (38–43). However, even these pioneering studies have not yet systematically explored the interfaces regulating SC assembly.

Here we present genetic evidence for direct interactions between specific residues in SYP-1, -3 and -4 in *C. elegans.* We leveraged a previously identified temperature-sensitive mutation, *syp-1^K42E^* (24), to perform a saturating forward suppressor screen. All suppressor mutations mapped to three short stretches in SC proteins, which likely represent a protein-protein interaction interface. Beyond their location within the middle of the SC, several lines of evidence support direct physical interaction. First, mutations were in conserved residues. Second, most mutations altered charged residues, compensating for the charge swap in the original mutation. Third, the suppressors restored not only the fertility defect that formed the basis of the screen but also other aspects of SC dysfunction like incomplete SC assembly, SC instability and weaker crossover interference. Fourth, the degree of suppression correlated with the nature of the mutation, with charge-neutral mutations conferring the weakest suppression. The extensive coverage in our screen further suggests that we isolated most of the possible suppressors and that other SC proteins are unlikely to participate directly in this interface.

The simplest model to explain our findings is a charge-charge interaction interface involving the three proteins, SYP-1, -3 and -4. Such a model readily accounts for the suppression of the original charge-swap mutation in SYP-1 position 42 by compensatory charge mutations in SYP-1, -3 and -4. The SYP-1/-3/-4 interface could in turn stabilize large-scale interactions between SC subunits to allow the SC to hold the axes of the two homologs together and/or to spread along chromosomes to synapse them end-to-end (44). However, without additional structural information, it would be challenging to test this model. Different model systems offer limited help due to the rapid sequence divergence of SC proteins (12–14). For instance, while the polymerization of the mammalian SC was proposed to involve interactions between tetramers of the human functional homolog of SYP-1, SYCP1 (39), these are mediated by hydrophobic residues, not charge-charge interactions as we propose here. Excitingly, the recent discovery of SKR-1/2 as a component of the SC in *C. elegans* allowed for the purification of recombinantly expressed worm SC (21). Future work in this system will potentially test the impact of the original and suppressor mutations on SC structure.

Recent studies in worms and plants demonstrated that the SC directly regulates crossover distribution (15, 45–49). The data in worms relied on perturbations to SC components, including partial depletion of SC proteins (15), the *syp-1^K42E^* mutant used here (24), and a serendipitous alteration in the C-terminus of SYP-4 (*syp-4^CmutFlag^(ie25);* (22)). These perturbations allow homologs to synapse and crossovers to form but reduce crossover interference so that each chromosome undergoes more than one crossover. Here, we observe multiple suppressors of the homolog synapsis function of the SC that also restore crossover interference. Strikingly, weak suppressors of the former conferred only partial suppression of the latter phenotype. Our findings, therefore, strengthen the idea that crossover interference is mediated by a mechanism that depends on the integrity of the SC (45, 46, 48–51). Destabilization of the SC by *syp-1^K42E^* (24) and restoration of stability by the suppressors (Figure 3G-H) further suggest that the biophysical properties of the SC control the strength of crossover interference.

Although the protein sequences of SC components are overall highly diverged in *Caenorhabditis* (12), the sites of the original and suppressor mutations tend to be well conserved. Sixteen out of 22 suppressor mutations occurred at sites that are 100% conserved in the Elegans and Japonica groups (Figure 4G), including the strongest suppressors we identified, SYP-3^D62V^ and SYP-4^E90K^. Similarly, we observed very little variation at suppressor sites in wild *Caenorhabditis* isolates. This conservation across and within species supports the importance of these positions for SC function.

Given the conservation of many of the suppressor sites, it is surprising that the suppressor mutations alone do not exhibit meiotic phenotypes (Figure 5). While the suppressor sites may not be important for SC function in the absence of *syp-1^K42E^*, their lack of phenotype is also consistent with a multivalent interaction interface. If SC assembly relies on multiple points of contact among its subunits (each likely composed of multiple interacting residues), altering one such interaction interface may not have a discernable effect. Consistent with this idea, combining two suppressors exposed a mild fertility defect (Figure 5C). Nonetheless, redundancy provides a poor mechanism to account for the purifying selection these residues seem to be subjected to, and it does not provide a good explanation for the strong phenotype exhibited by the single substitution in *syp-1^K42E^*. Alternatively, the suppressor sites may play a more important role in natural populations that are exposed to many biotic and abiotic stressors that sensitize meiotic fitness, and where minor fertility defects play out over many generations. We have tested one such stressor – temperature extreme – and did not observe a significant effect. However, we cannot rule out minor effects below our detection level or a role for other environmental stressors.

The epistasis we observe between *syp-1^K42E^* and the suppressor mutations has implications for the evolutionary trajectories of SYP-1 in *Caenorhabditis*. The SYP-1 fitness landscape in the presence or absence of a suppressor mutation is distinctly different, with only the former allowing a subsequent *syp-1^K42E^* mutation. In this regard, the suppressor mutations that are not conserved (*e.g., syp-1^T35I^*) are particularly interesting as they change the mutational landscape that could be sampled by SYP-1 (52, 53).

Our work demonstrates the power of genetic analysis in general, and suppressor screens in particular, to illuminate molecular mechanisms in metazoans. Several factors assisted in the success of the screen: the temperature sensitivity of the original mutation which allowed us to grow a large number of worms at the permissive temperature; the near complete sterility at the non-permissive temperature, conferring very tight selection; and the use of ENU as a mutagen, rather the more commonly used EMS (54, 55), which likely helped in isolating substitutions, since EMS mostly generates premature stop codons (56, 57). In the future, similar screens may prove particularly important in probing protein assemblies that are refractory to biochemical studies, as well as for studying cellular structures that assemble through condensation, as is the case for the SC (31), due to their reliance on weak multivalent interactions (58).

## Materials and Methods

### Worm strains

All strains were grown according to standard methods (59) and maintained at 20°C, with the exception of *gfp::cosa-1 (meIs8) II; syp-1^K42E^(slc11) V* which was maintained at 15°C. For experiments carried out at 15°C or 25°C (Figure 4 and Figure 5), worms were shifted to the indicated temperature as L4s, 24 hours prior to analysis. See below for growth conditions for the suppressor screen.

The following strains were generated via CRISPR/Cas9 by injection into *gfp::cosa-1 (meIs8) II* worms (29): *gfp::cosa-1 (meIs8) II; syp-1^K42E^ (slc11) V* (original mutation, made previously in (24)), *syp-3^D62N^ (slc41) I; gfp::cosa-1 (meIs8) II, syp-3^D62V^ (slc40) V ; gfp::cosa-1 (meIs8) II,* and *syp-4^E90K^ (slc42) I; gfp::cosa-1 (meIs8) II*. To make the double suppressor worms, we injected a *syp-3* CRISPR mix into *syp-4^E90K^ (slc42) I; gfp::cosa-1 (meIs8) II*. To make the suppressed strain for live imaging, *HTP-3::wrmScarlet (slc1) syp-3 I*; ieSi11[EmGFP-SYP-3] II; *syp-1^T35I+K42E^(slc68) V,* we injected into *HTP-3::wrmScarlet (slc1) syp-3 I; ieSi11[EmGFP-SYP-3] II* with *syp-1* CRISPR mix. We made suppressed strains (*syp-3^D62N^ (slc41) I; gfp::cosa-1 (meIs8) II; syp-1^K42E^ (slc11) V, syp-3^D62V^ (slc40) V; gfp::cosa-1 (meIs8) II; syp-1^K42E^ (slc11) V,* and*; syp-4^E90K^ (slc42) I; gfp::cosa-1 (meIs8) II; syp-1^K42E^ (slc11) V,* by crossing worms containing only the suppressor mutation, generated via CRISPR, with *gfp::cosa-1 (meIs8) II; syp-1^K42E^ (slc11) V.* For a complete list of strains used in this study, see Table S4.

### Total progeny and percent males

We picked at least 10 individual L4s from each strain and grew them at the indicated temperature. We transferred the worms to a new plate every day for five days and counted the total number of adult progeny and percent males 3 – 5 days later, depending on the temperature. For progeny counts of the heterozygous suppressed strains, we crossed *gfp::cosa-1 (meIs8) II; syp-1^K42E^ (slc11) V* to either *syp-3^D62V^ (slc40) I*; *gfp::cosa-1 (meIs8) II; syp-1^K42E^ (slc11) V*, *syp-3^D62N^ (slc41) I*; *gfp::cosa-1 (meIs8) II; syp-1^K42E^ (slc11) V*, or *syp-4^E90K^ (slc42) I*; *gfp::cosa-1 (meIs8) II; syp-1^K42E^ (slc11) V*. We selected at least 10 heterozygous F1 animals from each cross and performed the progeny count experiment as described. After the last day of counts, we genotyped the F1 to ensure it was a heterozygote.

### Mutagenesis with N-ethyl-N-nitrosourea (ENU) and screen for suppressors

We grew three 10 cm plates of mixed-age population *gfp::cosa-1; syp-1^K42E^* at 15°C on 8P media with NA22 bacteria until the plates were dense with worms but not starved. We washed the worms off of the plates and bleached them to generate a large number of synchronous embryos. We used these synchronized worms for mutagenesis when most worms were at the L4 developmental stage (four days after bleaching at 15° C). We washed the synchronized L4s once with M9 + 0.01% TritonX, resuspended them in five ml M9 + 0.01% TritonX in a 50 ml tube, and added 50 ul of 50 mM ENU. The worms in ENU were rocked gently at room temperature for four hours. After mutagenesis, we washed the worms three times in 30 ml M9 + 0.01% TritonX, allowing the worms to settle between each wash. We then distributed the worms (P^0^ generation), approximately 500 worms per plate, among 100 plates (10 cm 8P media plates with NA22 bacteria), and let them recover for five days at 15° C.

After five days at 15° C, when many of the F^1^ animals had reached the L4 developmental stage, we moved the plates to 25° C. We fed each plate 1ml, 100X concentrated overnight culture of NA22 three times, after 24, 48 and 72 hours at 25°C. On the 4^th^ day after the shift to 25° C (F^2^ animals should be young adults), we chunked all plates (∼2 x 2 cm chunk). We then chunked plates as needed to prevent immediate starvation (up to three times), and monitored for the ability of the worms to proliferate at 25°C.

### Notes on suppressed strains isolated from the screen

Four of the suppressed genotypes characterized in our experiments (Figure 3) were isolated as homozygous mutants from the screen: *gfp::cosa-1 (meIs8) II; syp-1^K42E+T35I^(slc11 + slc60) V*, *gfp::cosa-1 (meIs8) II; syp-1^K42E+V39I^ (slc11 + slc65) V, gfp::cosa-1 (meIs8); syp-1^K42E+S24L^ (slc11 + slc49) V,* and *gfp::cosa-1 (meIs8) II; syp-1^K42E^ (slc11); syp-4^A81T^ (slc63).* Our whole genome sequencing revealed that these strains were unlikely to have phenotypes caused by mutations in other genes expressed during meiosis. Other mutations in meiotic expressed genes in *gfp::cosa-1 (meIs8) II; syp-1^K42E+T35I^(slc11 + slc60) V* and *gfp::cosa-1 (meIs8) II; syp-1^K42E+V39I^ (slc11 + slc65) V* were predicted to be low impact (synonymous, upstream or downstream variants, Table S4). *gfp::cosa-1 (meIs8); syp-1^K42E+S24L^ (slc11 + slc49) V* has a missense mutation in ZK430.5, which is related to *C. elegans* separase (*sep-1*). However, RNAi of ZK430.5 has no phenotype (60). *gfp::cosa-1 (meIs8) II; syp-1^K42E^ (slc11); syp-4^A81T^ (slc63)* has a missense mutation in C30G12.6 which is predicted to be a MPK-1 substrate. However, RNAi phenotypes due to C30G12.6 knockdown were only apparent in a sensitized *mpk-1* background (61). Therefore, we attribute all meiotic phenotypes to the mutations in SC proteins. All other suppressors were re-engineered into the parental *gfp::cosa-1 (meIs8) II; syp-1^K42E^ (slc11) V* strain as described above (Table S4).

### Whole genome sequencing

To extract genomic DNA for whole genome sequencing, we grew five, 6cm plates of each suppressed strain until they were just starved. We washed the worms off plates and into a 15ml conical tube with M9. We performed four additional washes (three quick washes of ∼ 1 minute and one long, 30 min wash) to remove as much bacteria as possible, collecting the worms by gentle centrifugation between each wash so that the worms formed a pellet but the bacteria remained floating in solution. After the last M9 wash, we pelleted the worms by centrifugation and removed as much buffer as possible. We then followed the Solid Tissues protocol from Zymo Quick-DNA Miniprep Plus Kit (cat#D4068) with slight modifications to improve the 260/230 ratio of the eluted DNA. We found that it was important to add a dry spin prior to DNA elution to prevent elevated absorbance at 230nm wavelength. We eluted in 50ul DNA elution buffer (10mM Tris-HCL, 0.1mM EDTA).

For library preparation, we used the Nextera DNA Flex library prep kit which includes minimal PCR (4 cycles). Samples were sequenced on an Illumina NovaSeq on a 2 x 150 bp run to sequencing depth of 30X coverage. Library preparation and whole genome sequencing were carried out at the High Throughput Genomics facility at the Huntsman Cancer Institute.

### Genome assembly and variant calling

We used BWA MEM (62) to align reads to the *C. elegans* reference genome (version WBcel235 from wormbase.org). We used GATK MarkDuplicates (Picard) to remove duplicate reads and GATK UnifiedGenotyper to call single nucleotide variants and indels relative to the reference genome (63, 64). We subtracted all variants present in the parental, non-ENU-treated strain *gfp::cosa-1 (meIs8) II; syp-1^K42E^ (slc11) V*, from each suppressed strain using Bedtools (65), creating a list of ENU induced mutations. We filtered the ENU induced mutations for homozygous variants that had a read depts of greater than 15 and a quality score greater than or equal to 200 using SnpSift (66). We annotated each of the remaining variants on the *C. elegans* genome and predicted its impact with SnpEff (67)

In total, we sequenced and analyzed 16 suppressed strains, *gfp::cosa-1 (meIs8) II,* and *gfp::cosa-1 (meIs8) II; syp-1^K42E^ (slc11) V*. Of the 16 suppressed strains, four contained candidate suppressor mutations in *syp-1* (*syp-1^S24L^*[twice], *syp-1^T35I^,* and *syp-1^V39I^*) that we had already identified via Sanger sequencing. Among the suppressed strains, we found two pairs of genomes that contained the same suppressor mutations but shared blocks of ENU-induced SNPs; one pair containing the *syp-4^E90G^*mutation (genomes 16190X1 and 16190X2) and the other containing the *syp-4^E90K^* mutation (genomes 16190X7 and 16191X3). This is contrast to all other genomes, which each had a completely unique mutational profile. We hypothesize that these shared blocks of mutations stem from mutagenic events in mitotic germline nuclei in the P^0^s, although their exact source is unknown. Since we could not verify the independence of the mutations in genomes 16190X1/16190X2 and 16190X7/16191X3, we only included one genome from each pair in all subsequent analyses. As a result, statistics surrounding ENU induced mutations were calculated from the 14 suppressed genomes that had unique mutational profiles. All genome sequences generated in this study are available at www.ncbi.nlm.nih.gov under BioProject PRJNA1006752.

### Coiled-coil predictions

We used paircoil2 (68) to predict coiled-coil domains and to annotate heptad repeats in Figure 3. The standard deviation of coiled-coil scores in Figure S5 was calculated from *Caenorhabditis* species coiled-coil scores published previously (12).

### CRISPR genome editing

We generated an RNA mix by combining 4ul 200uM tracrRNA (IDT Alt-R® CRISPR-Cas9 tracrRNA), 3ul 200uM crRNA matching our gene of interest (IDT Alt-R® CRISPR-Cas9 crRNA) and 1ul *dpy-10* crRNA. We heated the RNA mix at 95°C for 5 minutes and then let it cool on the benchtop for 5 minutes. We next made an injection mix by combining 3.5ul of the RNA mix with 1ul Cas9 (IDT, Alt-R™ S.p. Cas9 Nuclease V3, 10ug/ul) 0.5ul 200uM *dpy-10* repair template, and 3ul repair template to our gene of interest (200uM). We injected 24hr post-L4 animals (P^0^s) and screened for Dpy and Rol F^1^s four to five days following injection. We singled Dpy and Rol F^1^s and genotyped them by extracting DNA from their progeny, performing PCR and a digest to distinguish the desired CRISPR allele from a wildtype allele. All crRNAs, repair templates and oligos used for genotyping are listed in Table S5.

### Immunofluorescence

Immunofluorescence was carried out essentially as previously described (69).We dissected 24hr post-L4 animals in 30ul Egg Buffer with 0.01% Tween-20 and 0.005% tetramisole (69). We added equal volume of a 2% formaldehyde solution in 1× Egg Buffer and incubated for 1 min in order to fix the samples. We transferred dissected samples to a HistoBond microscope slide (VWR) and froze samples on dry ice. We quickly removed the coverslip and immersed samples in –20°C methanol for 1 min. We washed samples 3 × 5 min in PBST (0.1% Tween-20), blocked the sample in Roche Block (product no. 11096176001) and incubated in primary antibodies overnight at 4°C in Roche Block. Following primary antibody incubation, we washed samples 3 × 5 min in PBST and incubated in secondary antibodies for 2 hr at room temperature. Finally, we washed samples for 10 min in PBST, 10 min in DAPI (5 μg/μl), and 10 min in PBST. We mounted samples in NPG-glycerol.

Primary antibody concentrations used were as follows: rabbit anti-SYP-2 1:500 (17), guinea pig anti-HTP-3 1:500 (70), mouse anti-GFP 1:1000 (Sigma Aldrich, #118144600011). Secondary antibody concentrations used were as follows: donkey anti-rabbit Alexa Fluor 647 1:500 (Jackson laboratories #711-605-152), donkey anti-guinea pig Cy3 1:500 (Jackson laboratories #706-165-148), donkey anti-mouse Alexa Fluor 488 1:500 (Jackson laboratories #715-545-150).

### Quantification of GFP::COSA-1 foci and asynapsis

We counted bright GFP::COSA-1 foci in orthogonal projections of confocal images with DAPI staining and HTP-3, SYP-2 and GFP antibody staining. We counted foci in non-overlapping single nuclei from the mid- and late-pachytene region of the gonad. To measure asynapsis, evaluated nuclei in early pachytene. We counted asynapsis as any region of HTP-3 staining with no SYP-2 staining.

### Measuring the transition zone

We generated whole gonad images by stitching together multiple (usually three) confocal image projections of DAPI-stained gonad segments. We identified the transition zone as the region of the gonad with crescent-shaped nuclear morphology. We defined the end of the transition zone as the last row of nuclei with at least two crescent-shaped nuclei followed by two or more rows with one or zero crescent-shaped nuclei. We defined the pachytene region of the gonad as starting at the transition zone and ending just before the nuclei become single file. We measured the length of both of these regions in Fiji (71) and divided the length of the transition zone by the length of the meiotic gonad.

### Statistical tests

In Figure 1B, we compared total progeny between *gfp::cosa-1; syp-1+* and *gfp::cosa-1; syp-1^K42E^* at 15°C and 25°C using unpaired t-tests. We used an ordinary one-way ANOVA to test for significant differences in total progeny (Figure 3A, Figure 5A-C, top), transition zone length (Figure 3F) and SC recruitment to chromosomes (Figure 3H) between *gfp::cosa-1; syp-1+, gfp::cosa-1; syp-1^K42E^* and suppressed strains. We used a Brown-Forsythe and Welch ANOVA test (for samples with unequal variances) to test for differences in percent males (Figure 3B, Figure 5A-C, bottom) and we used a Kruskal-Wallis test to test for differences in GFP::COSA-1 foci counts (Figure 3I, Figure 5D). In Figure 3, all comparisons are to *gfp::cosa-1; syp-1^K42E^* (black asterisks) and in Figure 5, all comparisons are to *gfp::cosa-1* (blue asterisks). Asterisks are representative of p-values as follows: * < 0.05, ** < 0.01, *** < 0.001, **** < 0.0001.

### Testing SC behavior at 26.5°C

We picked L4 animals from *syp-4^E90K^ (slc42) I; gfp::cosa-1 (meIs8) II;* and *gfp::cosa-1 (meIs8) II,* and grew them at 20°C for eight hours. We then moved both strains to 26.5°C for 16 hours. After 16 hours at 26.5°C (24 hours total post-L4), we dissected the worms and performed immunofluorescence.

### Live imaging

We dissected 24 hr post-L4 animals grown at 20°C on a coverslip in modified embryonic culture medium ((72), 84% Leibowitz L-15 medium, 9.3% Fetal calf serum, 1% 2 mM EGTA). We transferred dissected animals to a 2% agarose pad on a glass slide and sealed the coverslip with valap sealant (equal parts petroleum jelly, lanolin and paraffin). We imaged intact gonads within 20 minutes of dissecting on a Zeiss LSM880 confocal microscope with Airyscan and a 63 × 1.4 NA oil objective. We took z-stacks (step size 0.4um) that spanned a depth of approximately one nucleus (∼15 slices). All images took around 30 seconds to collect. The power for the 488 argon laser and the 561 nm diode laser was kept constant at 2.6 for all experiments and genotypes.

### Quantification of SC recruitment to chromosomes

To avoid artifacts of fixation and antibody staining, we performed SC recruitment measurements in live gonads. Since all SC components in worms are co-dependent for SC assembly, GFP::SYP-3 localization is a reliable proxy for SC stability (73). We used Imaris x64 9.7.2 (Oxford Instruments) to process and quantify all live imaging data. We cropped individual nuclei and used the signal from the axis protein HTP-3::wrmScarlet to define chromosomes using the Surface generator with default thresholds set by the absolute intensity of HTP-3::wrmScarlet signal. We measured the total SYP-3::GFP signal within the chromosomal surface and divided by the total SYP-3::GFP signal in the cropped nucleus to determine the percent SC localized to chromosomes.

## Acknowledgments

We would like to thank Sean Shadle for help with bioinformatics, Erik Jorgensen and Matt Labella for advice on using ENU, Yumi Kim for antibodies, Sara Nakielny for comments on the manuscript and editorial work, the scientific illustrator Maria Diaz de la Loza for graphical work and members of the Rog lab for discussions. Some worm strains were provided by the CGC, which is funded by NIH Office of Research Infrastructure Programs (P40OD010440). Research reported in this publication utilized the High-Throughput Genomics and Cancer Bioinformatics Shared Resource at Huntsman Cancer Institute at the University of Utah and was supported by the National Cancer Institute of the NIH (P30CA042014). LEK was supported by the Developmental Biology Training Grant from NICHD (T32HD007491). OR wishes to thank the Taft-Nicholson Center for Environmental Humanities Education Center for a Summer Fellowship. JEAM was supported by a Biology Research Scholar Award. This work was funded by NIGMS grant R35GM128804.

