## Supplemental Figures for "A suppressor screen *in C. elegans* identifies a multi-protein interaction interface that stabilizes the synaptonemal complex"

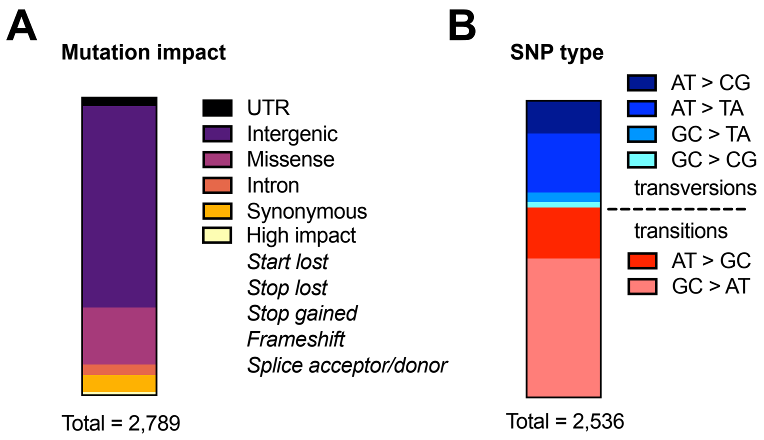


**Figure S1. Distribution of ENU caused mutations.** (A) Bar graph displaying the proportion of mutations affecting gene features. (B) Bar graph displaying proportion of each type of transition or transversion. Data in (A) and (B) is from 14 independent suppressed strains.

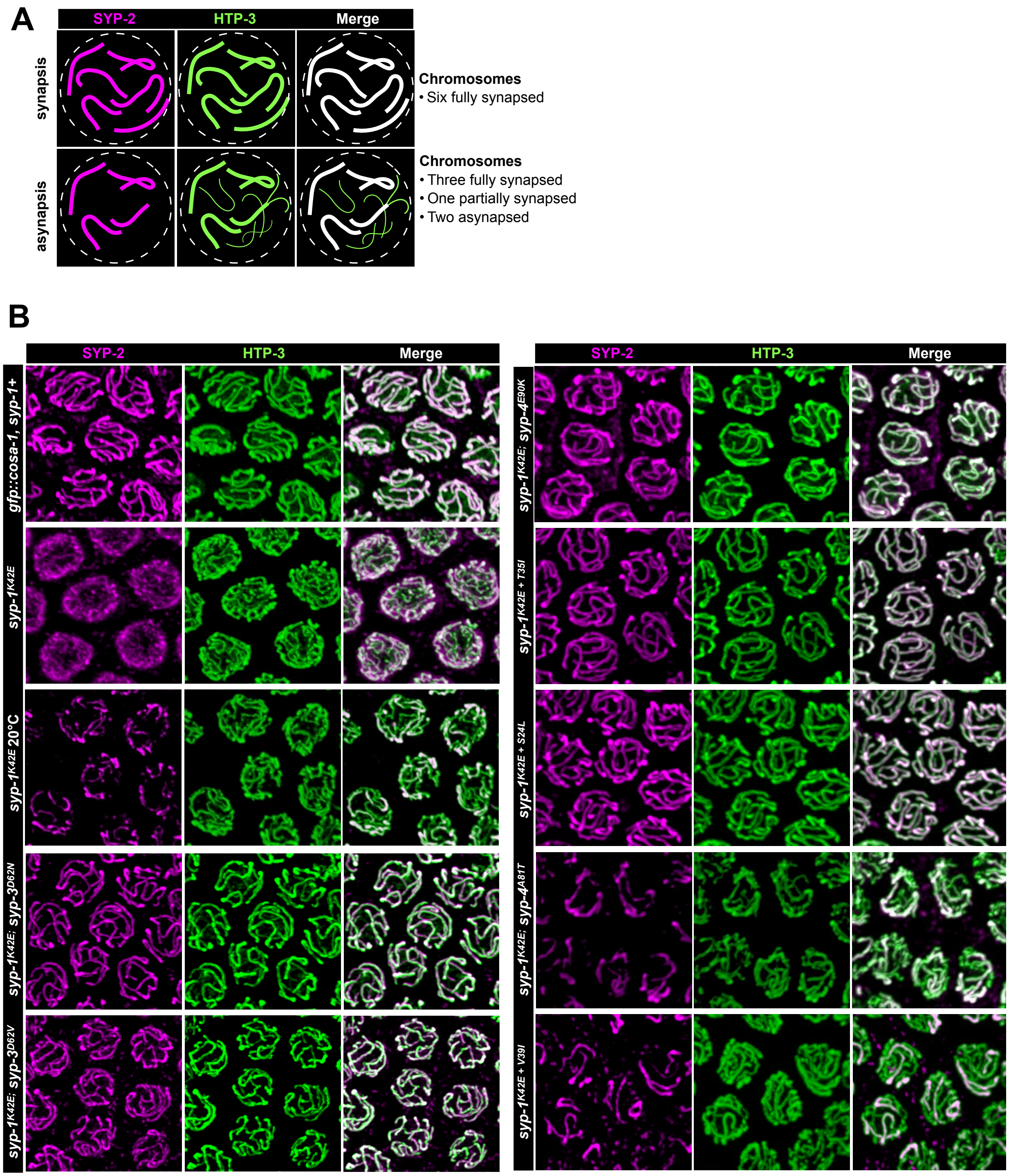


**Figure S2: Quantification of synapsis in suppressed strains.** (A) Interpretive diagram demonstrating complete synapsis and asynapsis, which includes partially synapsed chromosomes. Asynapsis can be identified in immunofluorescence images as axis staining (green) without SC (magenta). (B) Immunofluorescence images from Figure 4 with separated channels. All images are from animals grown for 24 hours at 25°C except for *syp-1*$^{K42E}$ 20°C. Asynapsis is apparent in *syp-1*$^{K42E}$ 20°C, *syp-1*$^{K42E}$*; syp-4*$^{A81T}$, and *syp-1*$^{K42E + V39I}$

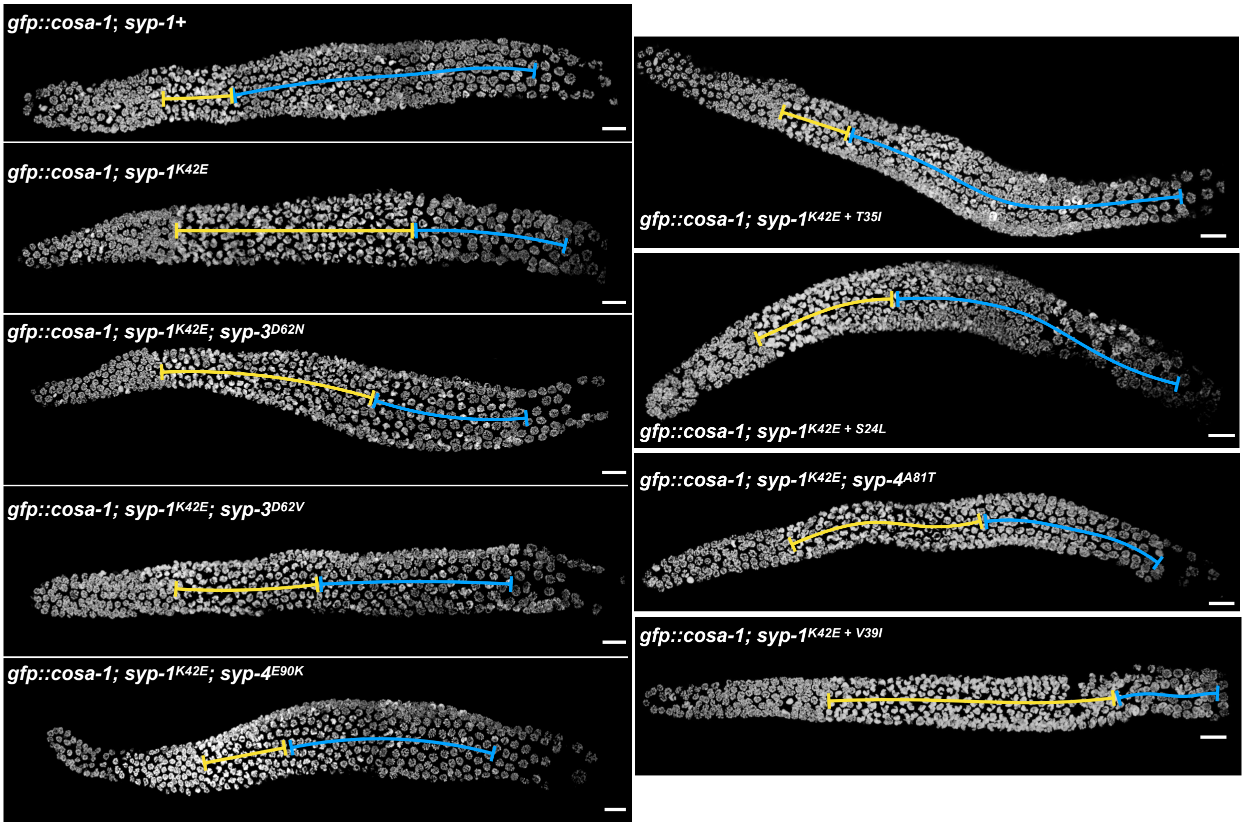


**Figure S3: Example transition zone images from suppressed strains and controls.** DAPI stained images of whole gonads from the indicated genotypes at 25°C. The transition zone, region of crescent-shaped nuclei, is labeled with a yellow line and the rest of meiosis is labeled with a blue line. Scale bars = 10μm.

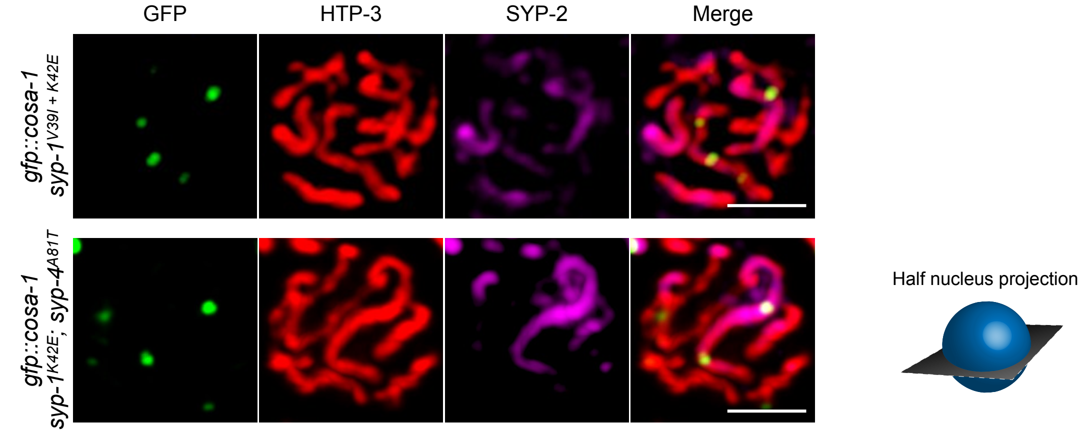


**Figure S4: Examples of chromosomes with multiple crossovers.** Half nucleus projections of immunofluorescence images from two suppressed strains: *syp-1*$^{V39I + K42E}$ and *syp-1*$^{K42E}$; *syp-4*$^{A81T}$. Scale bar = 2μm.

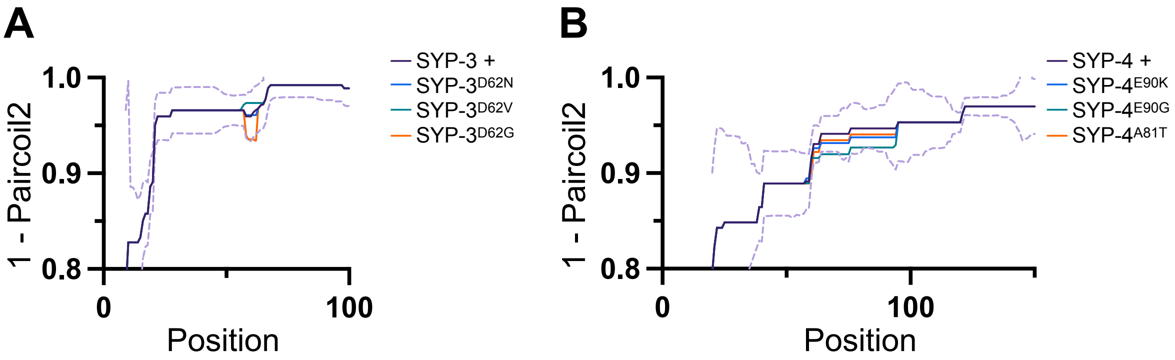


**Figure S5. Suppressor mutations do not significantly alter the predicted coiled-coil structure of SYP-3 and SYP-4.** Plots of coiled-coil scores per position for SYP-3 (A) and SYP-4 (B) wildtype and suppressor sequences. Both plots focus on the ~100 amino acids surrounding the suppressor mutation; the coiled-coil score for the rest of the protein is unchanged. The purple dashed line indicates one standard deviation of the coiled-coil score calculated from the scores across *Caenorhabditis*.

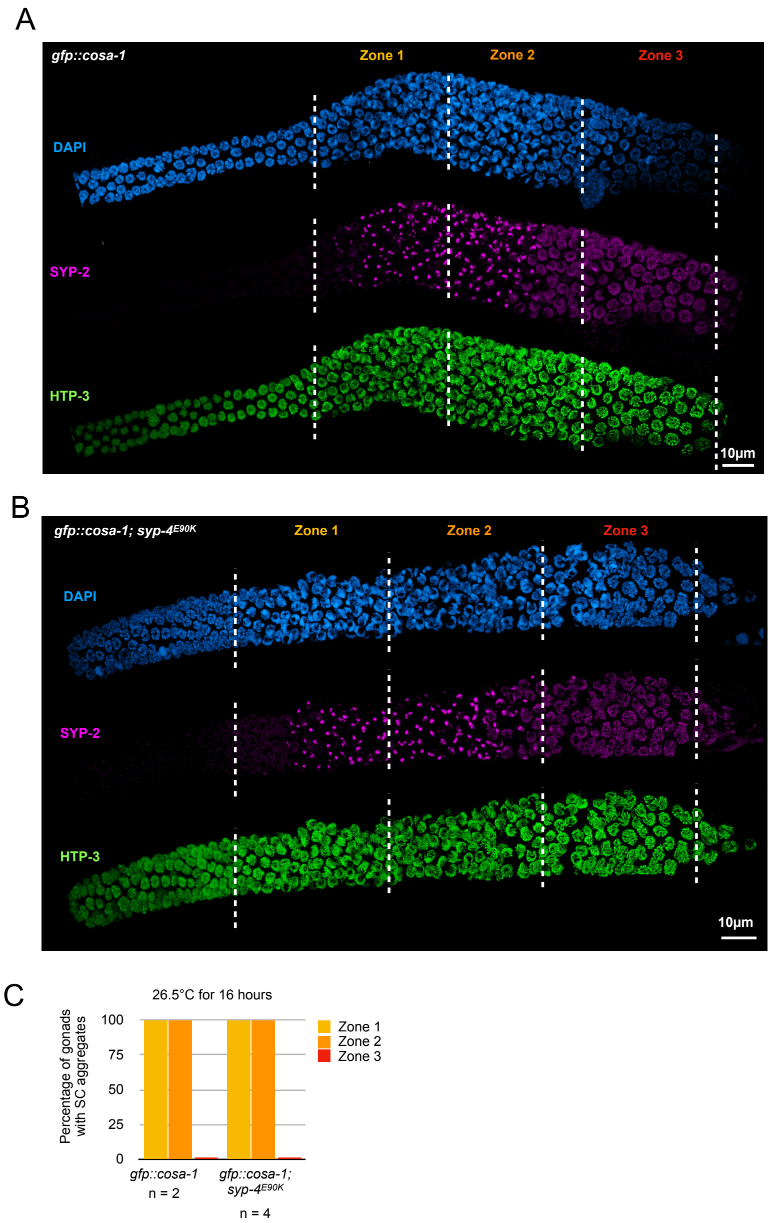


**Figure S6: Temperature extremes do not impact SC behavior in *gfp::cosa-1; syp-4$^{E90K}$*.** Representative whole gonad immunofluorescence images of *gfp::cosa-1* (A) and *gfp::cosa-1; syp-4$^{E90K}$* (B) after exposure to 26.5°C for 16 hours. The meiotic region of the gonad is divided into three zones of equal length. (C) Quantification of SC aggregates in each zone in each genotype after 16 hours at 26.5°C. We did not detect SC aggregates in zone three in any gonad of either genotype, presumably because SC that is already assembled is protected from aggregation at 26.5°C.
